# chTOG is a conserved mitotic error correction factor

**DOI:** 10.1101/2020.08.03.235325

**Authors:** Jacob A. Herman, Matthew P. Miller, Sue Biggins

## Abstract

Accurate chromosome segregation requires kinetochores on duplicated chromatids to biorient by attaching to dynamic microtubules from opposite spindle poles, which exerts forces to bring kinetochores under tension. However, kinetochores initially bind to MTs indiscriminately, resulting in errors that must be corrected. While the Aurora B protein kinase destabilizes low-tension attachments by phosphorylating kinetochores, low-tension attachments are intrinsically less stable than those under higher tension *in vitro* independent of Aurora activity. Intrinsic tension-sensitive behavior requires the microtubule regulator Stu2 (budding yeast Dis1/XMAP215 ortholog), which we demonstrate here is likely a conserved function for the TOG protein family. The human TOG protein, chTOG, localizes to kinetochores independent of microtubules by interacting with Hec1. We identify a chTOG mutant that regulates microtubule dynamics but accumulates erroneous kinetochore-microtubule attachments that Aurora B fails to destabilize. Thus, TOG proteins confer a unique, intrinsic error correction activity to kinetochores that ensures accurate chromosome segregation.

## Introduction

Eukaryotic cell division requires the duplication and accurate segregation of up to hundreds of chromosomes. In most species, chromosome segregation is carried out by a conserved network of dozens of kinetochore factors that assemble into a megadalton protein complex on centromeres to link chromosomes and dynamic microtubule polymers (Hara & Fukagawa, 2018). During mitosis, the microtubule cytoskeleton is organized into a bipolar spindle such that each and every pair of duplicated sister chromosomes becomes bioriented (attached to microtubules anchored to opposite poles). In this state, coordinated depolymerization of all kinetochore bound microtubules results in the accurate segregation of every chromosome. However, the biorientation process is error-prone, as early in mitosis both the mitotic spindle and the chromosomes lack organization, yet kinetochores begin forming attachments (Cimini, Moree, Canman, & Salmon, 2003; Kapoor, Mayer, Coughlin, & Mitchison, 2000; Maiato, Gomes, Sousa, & Barisic, 2017). While some kinetochore pairs become properly bioriented, others attach to microtubules emanating from the same spindle pole and must be destabilized. It is well appreciated that tension generated by depolymerizing microtubules pulling across a pair of bioriented kinetochores is a key signal that attachments should be stabilized (Akiyoshi et al., 2010; King & Nicklas, 2000; Salmon & Bloom, 2017). This biochemical error correction system is primarily known to be regulated via the Aurora B protein kinase and its downstream targets, which specifically *destabilize* low-tension kinetochore-microtubule attachments (Biggins & Murray, 2001; Cheeseman, Chappie, Wilson-Kubalek, & Desai, 2006; J. G. DeLuca et al., 2006; D. Liu, Vader, Vromans, Lampson, & Lens, 2009). However, we recently demonstrated that the protein Stu2 (yeast member of the Dis1/XMAP215 family) confers tension-sensitive binding behaviors to reconstituted yeast kinetochore-microtubule attachments (Akiyoshi et al., 2010; Miller, Asbury, & Biggins, 2016). Moreover, this intrinsic tension-dependent activity functioned completely independent of Aurora B activity (Akiyoshi et al., 2010; London, Ceto, Ranish, & Biggins, 2012; Miller et al., 2016), suggesting that cells have multiple mechanisms to destabilize incorrect attachments.

Stu2 and the entire Dis1/XMAP215 family are well characterized microtubule regulators that contribute to the nucleation, polymerization, and organization of the cytoskeleton and spindle in both developing and somatic cells (Brouhard et al., 2008; Cullen, Deák, Glover, & Ohkura, 1999; Gard & Kirschner, 1987; Kosco et al., 2001; Milunović-Jevtić, Jevtić, Levy, & Gatlin, 2018; Roostalu, Cade, & Surrey, 2015; Shirasu-Hiza, Coughlin, & Mitchison, 2003). This protein family is thought to accomplish these diverse forms of microtubule regulation through two regulatory regions. First, these proteins contain an array of 2-5 TOG domains that are each capable of binding an α/β tubulin dimer. Second, an unstructured ‘basic patch’ enriched for Lysine and Arginine residues appears to contribute to a non-specific electrostatic interaction with the negatively charged microtubule lattice (Geyer, Miller, Brautigam, Biggins, & Rice, 2018; Wang & Huffaker, 1997; Widlund et al., 2011). *In vitro*, these two regulatory regions catalyze the nucleation and elongation of microtubule polymers. However, Stu2’s ability to confer tension dependent binding behavior to reconstituted yeast kinetochore-microtubule attachments appears to be independent of its role in regulating microtubules, as all measures of dynamicity remained unchanged when Stu2 was absent from reconstitutions (Miller et al., 2016, 2019). Thus far, reconstitution experiments have been the only means to specifically study Stu2/XMAP215 regulation of kinetochore-microtubule attachments as *in vivo* depletion studies result in dominant defects in organizing the mitotic spindle (Kosco et al., 2001; Miller et al., 2019). We recently described a Stu2 mutant that supported spindle formation in yeast cells, but not biorientation, which provided *in vivo* evidence that Stu2 functions as an error correction factor independent of its role organizing the mitotic spindle (Miller et al., 2019). However, this mutant does not function as a microtubule polymerase *in vitro* (Geyer et al., 2018), raising the possibility that these two activities are connected in cells.

Similarly, depletion of the human ortholog, chTOG (TOG/TOGp/CKAP5), results predominantly in multipolar spindle assembly defects (Cassimeris & Morabito, 2004; Gergely, Draviam, & Raff, 2003). Chromosome biorientation is possible with partial depletion, and these kinetochores exhibit dampened oscillations and decreased inter-kinetochore tension (Barr & Gergely, 2008; Cassimeris, Becker, & Carney, 2009). While these data suggest chTOG regulates kinetochore-microtubule attachments, it is not clear if its role is related to regulating microtubule dynamics. Separation of these activities in human cells has also been limited by the ability to express mutant chTOG proteins. These large proteins (225 kDa) are inefficient to transduce through chemical and viral means, and negatively affect proliferation when over expressed. Therefore, it has been assumed TOG proteins regulate kinetochore-microtubule attachments only by affecting microtubule polymerization rates and any mitotic error correction role this protein family performs in multicellular eukaryotes has remained unknown.

Here we demonstrate that the Stu2-dependent error correction process observed in budding yeast is a conserved process in human cells. Similar to the yeast proteins (Miller et al., 2016), we found that chTOG associates with and requires the conserved microtubule binding factor Hec1 for kinetochore localization. Additionally, we show that a pair of point mutations in chTOG’s ‘basic linker’ domain inhibits error correction activity but does not compromise its ability to regulate the microtubule cytoskeleton. Together, this work reveals that chTOG functions in an evolutionarily conserved manner to destabilize erroneous, low-tension attachments. Moreover, this activity is independent of Aurora B phospho-regulation of its key kinetochore substrate, Hec1/Ndc80. Our work further elucidates a largely uncharacterized, intrinsic mechanism by which kinetochores sense and respond to biomolecular forces in order to prevent errors in chromosome segregation.

## RESULTS

### A pool of chTOG resides at kinetochores, independent of microtubule plus ends

Previous microscopy studies suggested chTOG also localizes to kinetochores (Campbell, Amin, Varma, & Bidone, 2019; Gutierrez-Caballero, Burgess, Bayliss, & Royle, 2015), similar to the budding yeast ortholog (Miller et al., 2016, 2019); however, it was unclear if this population was simply bound to microtubule tips or was an artefact of exogenous protein over-expression. To address this, we used engineered HCT116 cells where endogenous chTOG alleles were epitope tagged with EGFP (Cherry et al., 2019) to determine whether chTOG specifically localizes to kinetochores throughout mitosis (Fig. 1a). We found that chTOG is largely excluded from the nucleus and did not robustly associate with anti-centromere antibody (ACA) until prometaphase when its kinetochore localization increased 2.5-fold as cells progressed through mitosis (Fig. 1a, b). To determine what fraction of this signal is specifically interacting with kinetochores and not microtubule tips, we treated cells with nocodazole to depolymerize microtubules. At least 60% of chTOG recruited to prometaphase kinetochores and 30% recruited to anaphase kinetochores is independent of microtubules (Fig. 1a, b). This trend was also observed in HeLa cells overexpressing exogenous EGFP-chTOG (Figure 1 – Figure Supplement 1a). To determine whether microtubule attachment delivers chTOG to kinetochores, we arrested cells in S phase with thymidine and released them into the cell cycle in the presence of nocodazole so that attachments never occurred. In this experiment, chTOG was still detected on kinetochores, consistent with a kinetochore-bound pool that is separate from microtubule tips (Fig. 1 a, b, Figure1 – Figure Supplement 1a).

**Figure 1.**
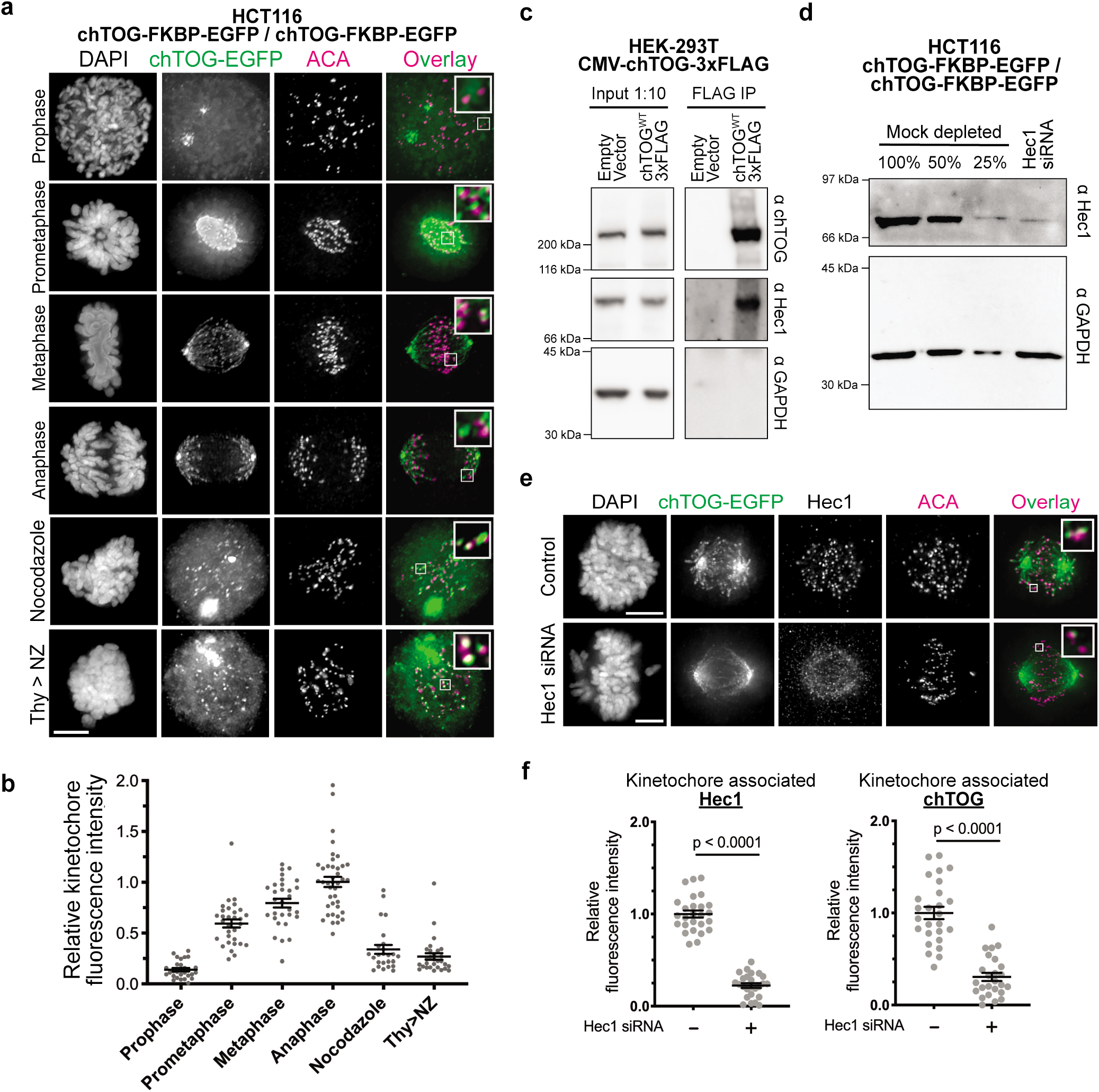
chTOG localizes to kinetochores during mitosis. (a) Immunofluorescence images of chTOG subcellular kinetochore proximal localization during mitosis, as visualized in HCT116 cells expressing endogenously epitope-tagged chTOG-EGFP. Anti-centromere protein antibody (ACA) staining marks the centromere binding proteins and representative images are shown with inlays of kinetochore proximal chTOG at each stage of mitosis, (b) Quantification of chTOG kinetochore association. Each data point represents mean chTOG-EGFP fluorescence intensity at all kinetochores in a single cell normalized to the mean value of the anaphase population, (c) 293T cells with either an empty vector control or overexpressed chTOG-3Flag were immunoprecipitated using anti-Flag antibody. Immunoblots of the input (left) or Flag IP (right) show that the endogenous Hec1/Ndc80 protein specifically co-purified with chTOG. Endogenous and epitope tagged chTOG cannot be individually resolved by anti-chTOG immunoblotting because the 3Flag tag increases the protein’s predicted MW by only 3%. Anti-GAPDH served as a non-specific control, (d) Immunoblotting with anti-Hed antibodies was performed on samples of mock-depleted lysate that were diluted to contain the indicated percent of total protein and compared to a lysate prepared from a population of HCT116 cells treated with Hec1 siRNA. Greater than 75% of Hec1 protein was depleted in the siRNA-treated cells. Anti-GAPDH is a loading control, (e) Kinetochore localization of Hec1 and chTOG in Hec1 depleted HCT116 cells was determined by fluorescence microscopy. Representative images are shown and were quantified in (f) to show both Hec1 antibody staining (left) and endogenously tagged chTOG-EGFP signal (right) at kinetochores decreased by ~80% in siRNA-treated HCT116 cells. Each data point represents mean fluorescence intensity at all kinetochores in a single cell normalized to the mean value of the mock depleted population. All scale bars are 5 μm; inlays have been background subtracted independently; all graphs display mean values, standard error of the mean, from three experimental replicates. Unpaired t test used to calculate p values.

chTOG and its budding yeast ortholog, Stu2, physically interact with the NDC80 kinetochore complex *in vitro* (Miller et al., 2016). To test whether they associate in human cells, we immuno-purified FLAG-tagged chTOG from HEK-293T cells under conditions refractory to microtubule formation and found that the endogenous Hec1 protein co-purifies (Fig. 1c). We therefore tested whether the kinetochore-bound pool of chTOG depends on Hec1 by depleting Hec1 from cells and quantifying the chTOG-EGFP intensity proximal to kinetochores (Figure 1d-f, Figure 1 – Figure Supplement 1b-d). Hec1 depletion ablates kinetochore-microtubule attachments (J. G. DeLuca et al., 2006), so we expected to see a disruption of the chTOG tipbound pool that would be similar to nocodazole treatment (40% reduction) and any further decrease would be due to defects in chTOG binding to kinetochores. We instead observed a 75% decrease in chTOG colocalization with kinetochores, consistent with the Ndc80 complex serving as the receptor for chTOG (Fig. 1e,f Figure 1 – Figure Supplement 1c,d). Therefore, a pool of chTOG localizes to kinetochores in a Hec1-dependent manner that is distinct from the chTOG at the microtubule plus-ends, suggesting a function for chTOG on kinetochores.

**Figure 1 – Figure Supplement 1.**
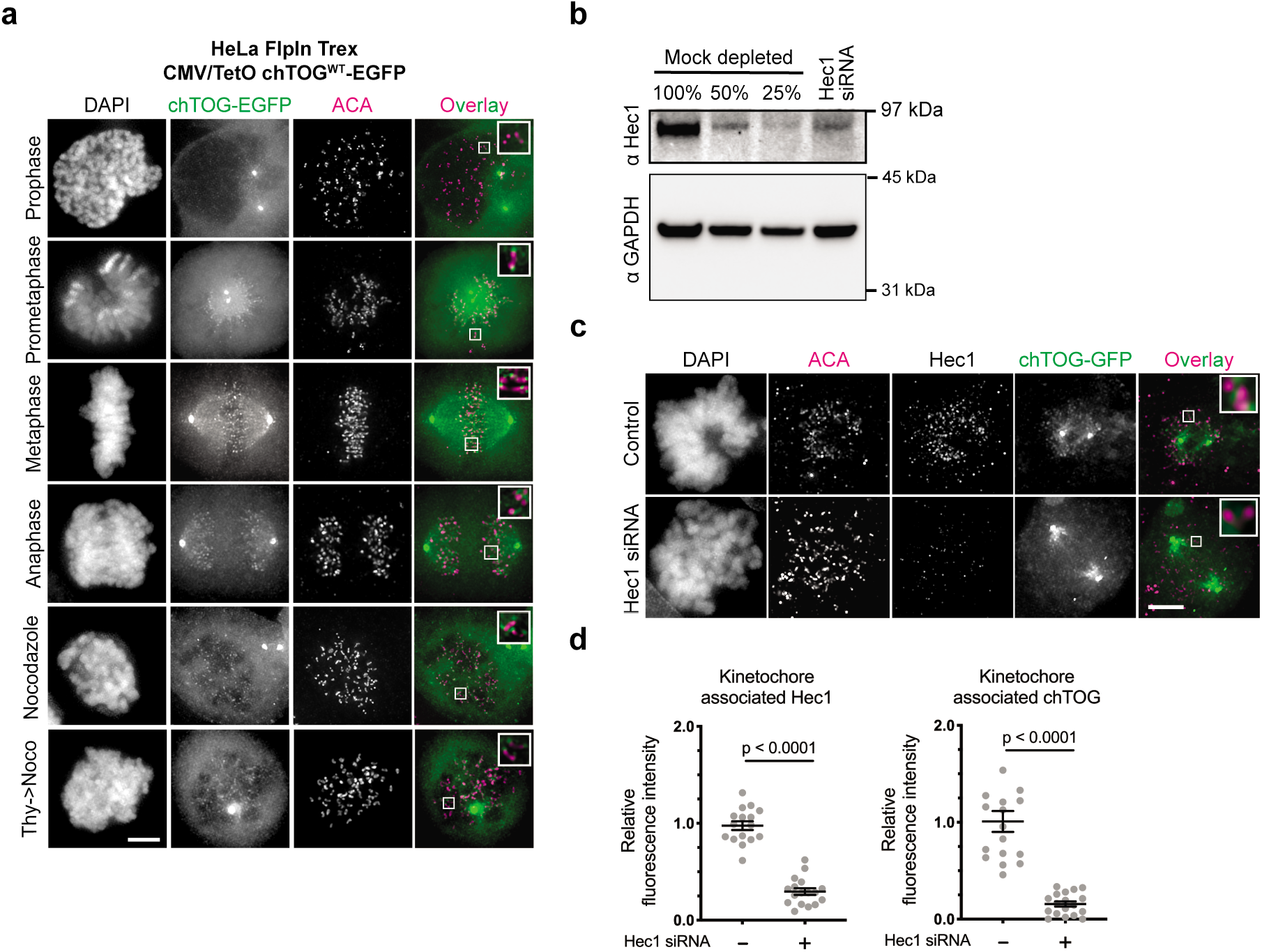
Exogenously expressed chTOG-EGFP localizes to kinetochores in HeLa cells. (a) Representative immunofluorescence images of chTOG subcellular kinetochore-proximal localization during mitosis, as visualized in HeLa cells where exogenous chTOG-EGFP was expressed by addition of 1 μg/mL doxycycline. Anti-centromere antibody (ACA) staining marks the centromere binding proteins and inlays of kinetochore proximal chTOG are shown in the boxes, (b) siRNA targeting Hed depleted more than 50% of the protein in a population of HeLa cells. Immunoblotting with anti-Hed antibodies was performed on samples of mock-depleted lysate that were diluted to contain the indicated percent of total protein and compared to a lysate prepared from a population of HeLa cells treated with Hed siRNA, (c) Representative images from immunofluorescence microscopy quantified in (d) showing Hed siRNA depletes 80% of the Hed and chTOG proteins from kinetochores in HeLa cells. Each data point represents the mean fluorescence intensity at all kinetochores in a single cell normalized to the mean value of the mock depleted population. All scale bars are 5 pm; inlays have been background subtracted independently; all graphs display mean values, standard error of the mean, from two experimental replicates. Unpaired t test used to calculate p values.

### Two residues in basic linker are essential for viability in yeast and human cells

To determine the role of chTOG at the kinetochore, we required a mutant that specifically inhibited its kinetochore function without affecting the protein’s numerous other microtubule-based activities. However, chTOG is an extremely large, multidomain protein consisting of 2032 residues, making it difficult to identify a separation of function mutant. It regulates microtubule dynamics using an array of five TOG domains (S Charrasse et al., 1998; Sophie Charrasse et al., 1995; Spittle, Charrasse, Larroque, & Cassimeris, 2000) (Fig. 2a). Additionally, chTOG contains a flexible ‘basic linker’ region that *in vitro* experiments suggest provides a non-specific, electrostatic interaction with the negatively charged microtubule lattice to facilitate diffusion to the plus-end (Geyer et al., 2018; Wang & Huffaker, 1997; Widlund et al., 2011). Finally, there is a small domain within the C-terminus of chTOG that serves as a protein interaction hub to mediate its various intracellular localization patterns (Gutierrez-Caballero et al., 2015; van der Vaart et al., 2011). All of these protein elements are present in the budding yeast ortholog Stu2, which contains just two TOG domains but homo-dimerizes through a coiled-coil region (CC) (Fig. 2a). We therefore took advantage of yeast genetic tools to identify mutants that potentially inactivate its kinetochore function. Previous cross-linking mass-spectrometry with yeast proteins revealed that both the Stu2 basic linker and C-terminus interact with the Ndc80 complex, but only the C-terminus was required for kinetochore association (Miller et al., 2019). This suggested the basic linker may instead interact with the Ndc80 complex to regulate kinetochore-microtubule attachments.

**Figure 2.**
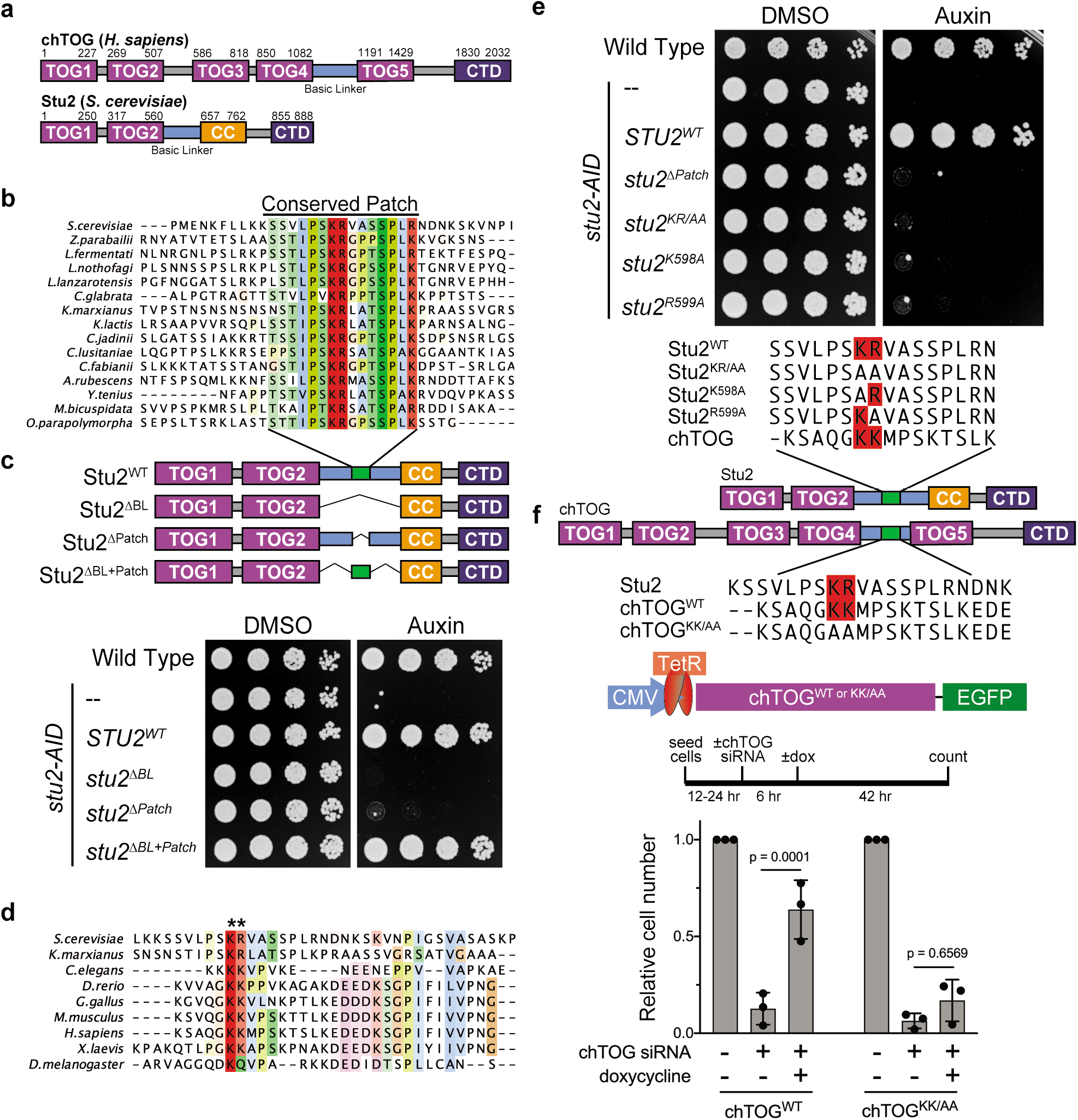
Two conserved basic patch residues are essential for Stu2/chTOG function. (a) Schematic of the yeast Stu2 and human chTOG proteins describing the domains in each protein. Specific residues marking domains are indicated on the top of each protein, (b) ClustalO multiple sequence alignment generated from full length S. *cerevisiae* Stu2 and related proteins in Ascomycota. Fifteen amino acids within the otherwise divergent −110 amino acid ‘basic linker’ are colored based on % conservation and biochemical properties of the side chain, (c) Schematic of the Stu2 mutant proteins used to investigate the essential nature of the conserved patch in S. *cerevisiae*. Cell viability was analyzed in stu2-AID strains expressing the indicated Stu2 mutant proteins by plating five-fold serial dilutions in the absence (left) and presence (right) of auxin to degrade the endogenous Stu2-AID protein, (d) ClustalO multiple sequence alignment generated from basic linker regions of metazoan and fungal species (colored similarly to (b)) highlights two conserved basic residues (asterisks), (e) The pair of conserved basic residues were mutated individually or as a pair to alanine (mutant sequences below) and found to be required for S. *cerevisiae* viability in a five-fold serial dilution growth assay when endogenous Stu2-AID is degraded (right, auxin), (f) Assay to analyze the ability of doxycycline-inducible codon-optimized, chTOG^wl^ and chTOG^KK/AA^ to support HeLa cell proliferation after siRNA-mediated depletion of endogenous chTOG. Relative cell counts were normalized to uninduced, mock depleted cells, mean values and standard deviation from three experiments displayed, p values determined with Tukey’s multiple comparisons test.

To identify a possible Ndc80 binding site in the basic linker, we aligned Stu2 orthologs from related Ascomycota and found that in addition to the previously described compositional positive charge bias, there was a conserved 15 amino acid sequence in the basic linker (Fig. 2b). To understand the role of this previously unidentified conserved patch, we ectopically expressed various *stu2* mutants under their native promoter in a strain where the endogenous allele was fused to an auxin-inducible degron (*stu2-AID*). We analyzed the viability of the various mutant cells in the presence of auxin, which degrades the endogenous Stu2-AID protein (Fig. 2c). These complementation studies demonstrated that the conserved patch was required for yeast viability (*stu2^ΔPatch^*). Strikingly, this patch (flanked by small flexible peptides) was sufficient to replace the entire 98 residue basic linker (*stu2^ΔBL+Patch^*), despite a 75% reduction in total length and an 83% reduction in positive residues (Fig. 2c). To determine if this patch is conserved from yeast to primates, we aligned the basic linkers among Ophistokonts and found that only two basic residues within the Stu2 essential patch are conserved outside Ascomycota (Fig. 2d). We therefore mutated Stu2 K598 and R599 to alanine as a pair (*stu2^KR/AA^*) or separately and found that despite normal mutant protein levels (data not shown), the loss of either residue was lethal to yeast (Fig. 2e). These data show that the two conserved residues, rather than the net charge of the basic linker, are essential for cell viability. Because the *in vitro* microtubule polymerase activity is regulated by net charge (Geyer et al., 2018; Widlund et al., 2011), our data suggest that there is a different essential role for the basic linker *in vivo*.

We set out to determine whether these conserved residues are required for chTOG function in human cells. Past studies of chTOG have primarily focused on its binding partners or RNAi depletion phenotypes due to its large size and multiple cellular functions (Gutierrez-Caballero et al., 2015; Hood et al., 2013). In addition, the constitutive overexpression of chTOG is toxic, making it difficult to study in mammalian cells. To overcome these technical challenges, we generated HeLa cell lines that harbor doxycycline-inducible, siRNA resistant alleles of chTOG-EGFP (chTOG^WT^) or chTOG[K1142, 1143A]-EGFP (chTOG^KK/AA^ or basic pair mutant) (Gossen & Bujard, 1992; O’Gorman, Fox, & Wahl, 1991; Taylor & McKeon, 1997). chTOG siRNA functioned in a dose-dependent manner allowing for partial (~60%) or near-complete (~90%) depletion (Figure 2 – Figure Supplement 1a). Doxycycline treatment of the 90% depleted cells induced expression of the ectopic chTOG proteins at equivalent levels, indicating that the mutations do not alter protein stability (Figure 2 – Figure Supplement 1a). We next tested whether the basic pair mutant affected chTOG localization. Using quantitative fluorescence microscopy in nocodazole-treated cells and biochemical analysis, we determined that the chTOG basic pair mutations did not affect its kinetochore association (Figure 2 – Figure Supplement 1b,c), which is consistent with our previous findings in budding yeast (Miller et al., 2019).

To determine whether the basic pair mutant could support cell viability, we analyzed cell proliferation in the depleted cells expressing the chTOG proteins. The WT chTOG protein restored viability, indicating that there were no off-target siRNA effects. In contrast, the chTOG basic pair mutant was not able to support cell proliferation after depletion of endogenous chTOG, despite expressing and localizing the same as WT chTOG (Fig. 2f). The failure to proliferate after chTOG depletion is due to a mitotic delay and the resulting chromosome segregation errors as previously documented (Cassimeris & Morabito, 2004; Gergely et al., 2003). To determine if chTOG^KK/AA^ induced similar defects, we stained the DNA and counted the mitotic index. After chTOG depletion, 20% of cells were mitotic, a marked increase from 5% of cells in a control population. Consistent with decreased cell viability, there was a similar mitotic delay in chTOG^KK/AA^ expressing cells (Figure 2 – Figure Supplement 1d). Taken together, these data suggest that there is an essential and conserved function of the basic linker in the chTOG protein that is unrelated to the compositional charge bias of this patch.

**Figure 2 – Figure Supplement 1.**
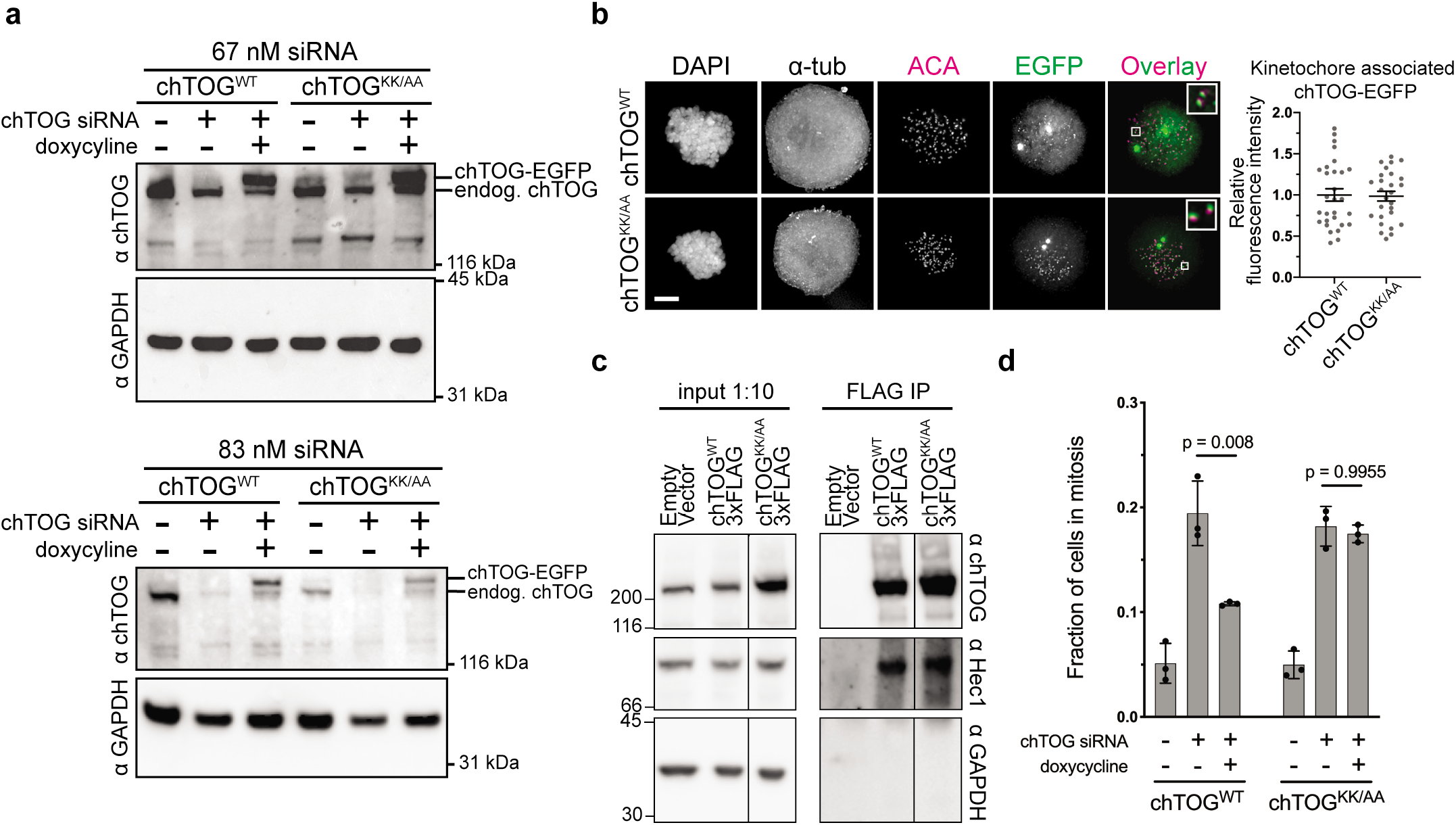
chTOG basic pair mutant protein levels and kinetochore localization are normal. (a) Immunoblots were performed on lysates of HeLa cells expressing EGFP epitope-tagged chTOG™ and chTOG^KK/AA^ proteins after depletion of the endogenous chTOG by siRNA with the indicated concentration of siRNA. All future experiments used 83 nM siRNA; GAPDH served as a loading control. Unlike 3FLAG epitope tag, both endogenous and EGFP tagged chTOG can be resolved with anti-chTOG antibodies (b) Representative images (left) and quantification (right) of chTOG™ and chTOG^KK/AA^ localization to kinetochores in the absence of endogenous chTOG. Cells were treated with nocodazole to eliminate microtubules. Each data point represents the mean chTOG-EGFP fluorescence intensity at all kinetochores in a single cell normalized to the mean value of chTOG™ expressing cells. Mean values and standard error of the mean from three experimental replicates shown, (c) chTOG was immunoprecipitated from 293T cells expressing either a control vector, or chTOGWT-3Flag or chTOG^KK/AA^-3FLAG proteins, using anti-Flag antibodies. Immunoblotting of the input lysates (left) and immunoprecipitations (right) show that the WT and mutant chTOG proteins co-purify endogenous Hed. Anti-GAPDH is shown as a non-specific control for the immunoprecipitation. This is the same immunoblot from Figure 1, and the line denotes where a single, non-relevant lane was cropped from the image, (d) Fraction of HeLa cells in mitosis after depletion of endogenous chTOG by siRNA and expression of EGFP epitope-tagged chTOG™ and chTOG^KK/AA^ proteins. Mean values and standard deviation from three experimental replicates shown.

### Mutating the basic pair does not alter dynamics or structure of the microtubule cytoskeleton

The mitotic arrest in cells expressing the basic pair mutant could arise from defects in a number of chTOG’s activities including: i) regulating cytoskeletal dynamics by nucleating and polymerizing microtubules, ii) organizing the mitotic spindle into a bipolar structure, or iii) regulating kinetochore-microtubule attachments. To understand if the basic linker contributed to these activities, we first analyzed the mutant by live cell TIRF microscopy on adherent interphase cells expressing EB1-mCherry (Tinevez et al., 2017; Tirnauer, Canman, Salmon, & Mitchison, 2002). chTOG requires functional TOG domains to bind microtubule plus-ends and elongate the polymer (Widlund et al., 2011), thus we verified that mutating the basic linker did not prevent the protein from localizing to growing plus-ends near EB1 (Fig. 3a). To determine whether microtubule assembly rates are affected, we next measured the velocity of EB1 comets after depletion of chTOG and observed an increase in microtubule assembly rates that was similarly suppressed by the expression of either chTOG^WT^ or chTOG^KK/AA^ in siRNA treated cells (Fig. 3b). Although this differs from other studies that reported either normal or decreased microtubule assembly rates upon chTOG knock down (Cassimeris et al., 2009; Ertych et al., 2014; van der Vaart et al., 2011), these discrepancies are likely due to technical differences (Supplementary note 1). Regardless, expression of the WT and basic pair mutant chTOG proteins restored microtubule dynamics equivalently (Fig. 3b), suggesting that the essential function of the basic pair mutant is not related to regulating microtubule dynamics. chTOG depletion leads to multipolar spindle formation (Cassimeris & Morabito, 2004; Gergely et al., 2003), so we next analyzed spindle morphology, and as expected, we found that nearly 50% of mitotic cells had multipolar spindles after chTOG depletion (Fig. 3c). However, expression of the WT and basic pair mutant proteins fully supported bipolar spindle formation when the endogenous chTOG protein was depleted (Fig. 3c). Thus, the basic pair mutant does not have detectable defects in the reported microtubule cytoskeleton functions of chTOG *in vivo* despite being essential for cell proliferation.

**Figure 3.**
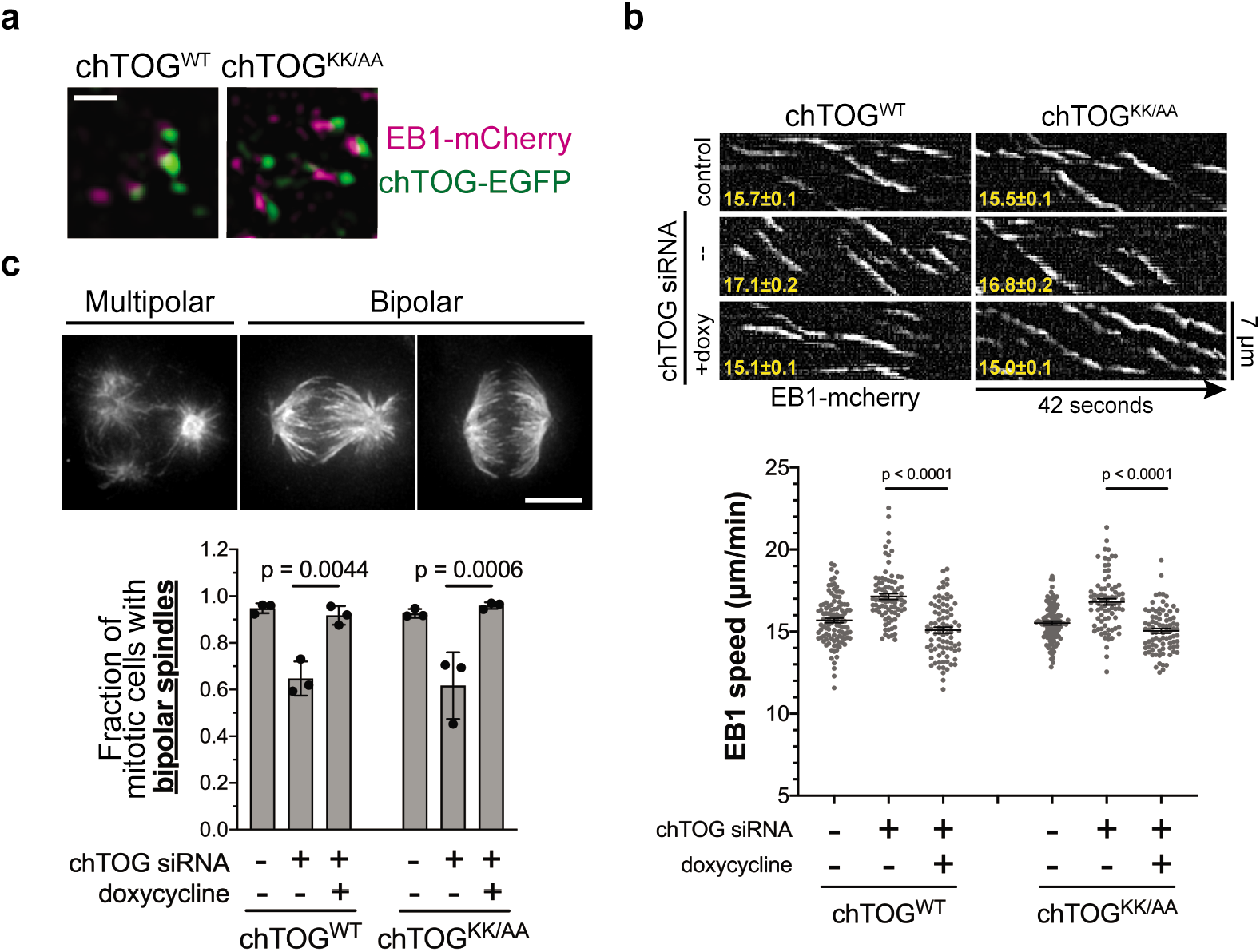
The chTOG basic pair mutant is proficient in microtubule regulation but exhibits defects in kinetochore-microtubule attachments. (a) Representative live-cell TIRF microscopy image of chTOG™ and chTOG^KK/AA^ binding to microtubule plus-ends as marked by EBl-mCherry. Images were isolated from movies used to generate kymographs of EB?-mCherry in chTOG™ and chTOG^KK/AA^ expressing cells in (b) with average EB1 track speed (μm/min) in bottom left and quantifications below. Each data point represents the mean EB1 track speed for each cell in a camera field. Mean values and standard error of the mean from three experimental replicates shown. Tukey’s multiple comparisons test used to calculate p values, (c) Representative images of each spindle phenotype observed in mitotic chTOG-depleted, chTOG™, or chTOG^KK/AA^ expressing cells while bipolar spindles exhibited two distinct phenotypes we first quantified the fraction of cells exhibiting multipolar or bipolar spindles. Mean values and standard deviation for three replicates reported, p values calculated with Tukey’s multiple comparisons test. All scale bars are 5 μm except for in (a) where scale bar is 1 μm.

### chTOG destabilizes incorrect kinetochore-microtubule attachments

Microtubule dynamics and spindle formation were not affected by mutating the basic pair; thus, we tested the possibility that chTOG functions like the yeast ortholog in regulating kinetochore-microtubule attachments. Further phenotypic analysis of mitotic cells expressing the basic pair mutant showed a defect in chromosome alignment, where 90% of cells formed a poorly organized metaphase plate (Fig. 4a). Most of the unaligned chromosomes were clustered at the poles with an excess of astral microtubules where they appeared to form stable syntelic or monotelic attachments (Fig. 4a, image inlays). These attachments were reminiscent of Aurora B kinase inhibition, suggesting that kinetochore-microtubule attachments were prematurely stabilized, allowing errors to persist and preventing chromosome alignment (Hauf et al., 2003; Kallio, McCleland, Todd Stukenberg, & Gorbsky, 2002). To better characterize the kinetochore-microtubule attachment state of chTOG^KK/AA^ expressing cells with unaligned chromosomes, we quantified the number of kinetochores with Mad1 staining because it specifically localizes to unattached kinetochores (Hoffman, Pearson, Yen, Howell, & Salmon, 2001; Howell et al., 2004). Because unperturbed prometaphase cells have many unaligned chromosomes, there is high error correction activity that destabilizes these erroneous attachments and generated an average of 36 Mad1 positive kinetochores (Fig. 4b). In contrast, an average of 13 kinetochores in chTOG depleted cells with unaligned chromosomes were Mad1 positive, which was rescued by expression of WT chTOG but not the basic pair mutant (Fig. 4b). These data suggest chTOG *destabilizes erroneous kinetochore-microtubule attachments*, and erroneous attachments persist when it is depleted or mutated. The persistence of these errors in cells expressing the basic pair mutant also explain the mitotic delay described above.

**Figure 4.**
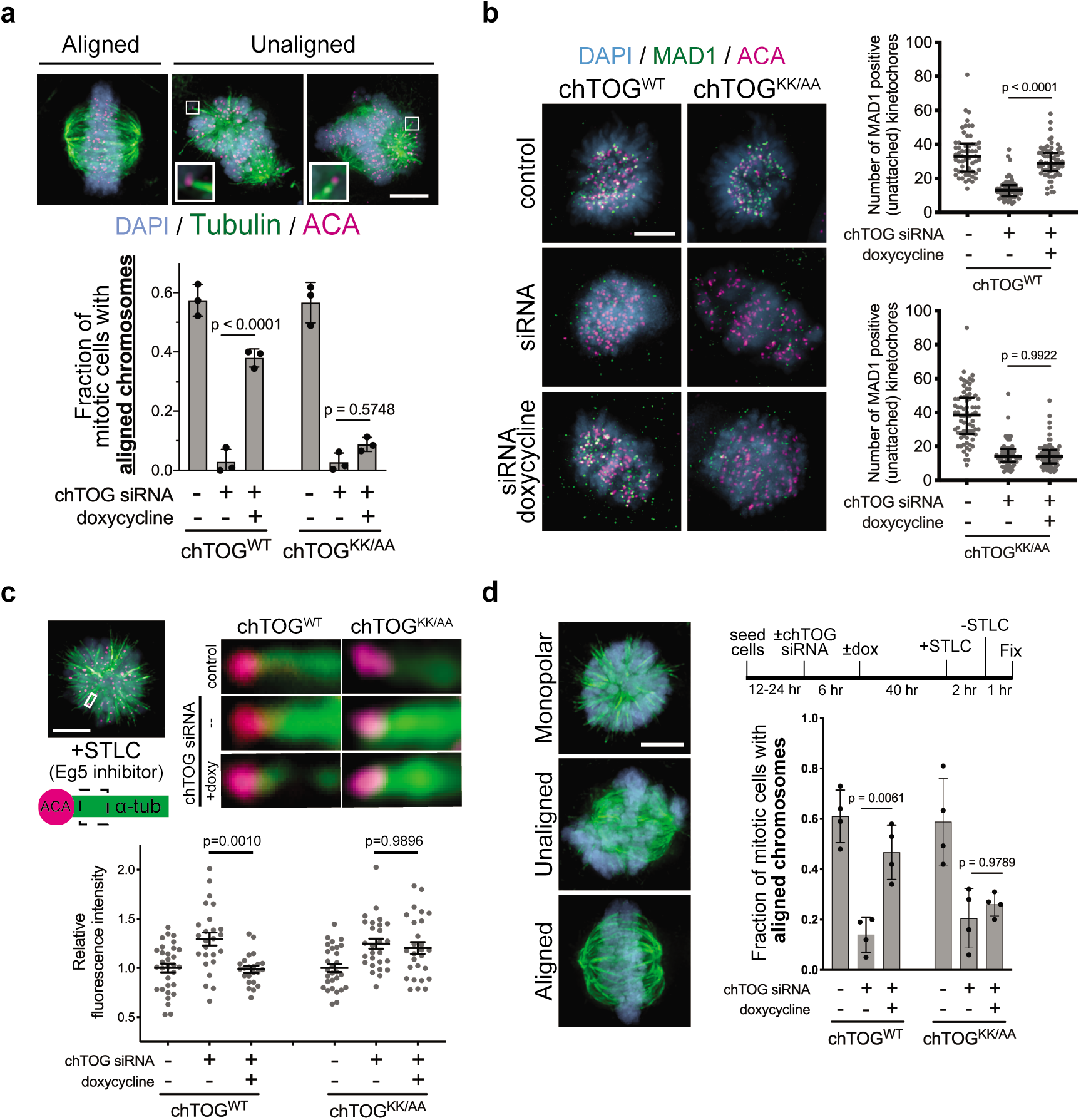
The chTOG basic pair mutant is defective in error correction. (a) Representative images of each chromosome alignment phenotype observed in mitotic chTOG-depleted, chTOG^WT^, or chTOG^KK/AA^ show a large fraction of chTOG^KK/AA^ expressing cells form bipolar spindles with excessive astral microtubules that attach to kinetochores (image inlays) and prevent chromosome alignment. Mean values and standard deviation for three replicates reported, (b) Representative images (left) and quantification (right) of Mad1 immunostaining as a marker for kinetochore-microtubule attachment state in chTOG depleted, chTOG^WT^ (top), or chTOG^KK/AA^ (bottom) expressing cells. Each data point represents the number of kinetochores with Mad1 puncta per cell, displaying median and interquartile ranges for three experimental replicates. Dunn’s multiple comparisons test used to calculate p values, (c) Monopolar spindles (top left) were formed by inhibiting Eg5/KIF11 with STLC to allow the fluorescence intensity quantification of kinetochore-bound microtubule bundles at low-tension, erroneous attachments in control cells or chTOG depleted cells expressing chTOG^WT^ or chTOG^KK/AA^ (right). Each data point is relative intensity normalized to the mean of uninduced, mock depleted cells. The mean and standard error of the mean for three experimental replicates are displayed, (d) Mitotic error correction was assayed by inducing errors with STLC to inhibit Eg5 and then washing out the inhibitor in control cells or chTOG depleted cells expressing chTOG^WT^ or chTOG^KK/AA^. The chromosome alignment phenotype (left) was quantified 60 minutes later; them mean and standard deviation of three experimental replicates are displayed. Unless otherwise stated, all p values were determined with Tukey’s multiple comparisons test. All scale bars are 5 μm; inlays have been background subtracted independently.

To further test the hypothesis that chTOG is required for error correction, we treated cells with a reversible Eg5/KIF11 inhibitor (STLC) to arrest them in a monopolar state, which enriches for low-tension erroneous attachments (Kapoor et al., 2000). We then assayed the relative number of microtubules bound by each kinetochore (Dudka et al., 2018). After chTOG depletion, kinetochore attached fibers contained 1.5x more microtubules per kinetochore (Fig. 4c). Expression of WT chTOG reversed this phenotype, while attachments remained hyper-stable when the basic pair mutant was expressed, even in the presence of endogenous chTOG (Figure 4 – Figure Supplement 1a). This provides further evidence that *chTOG turns over erroneous kinetochore-microtubule attachments* and that the basic pair is required for this activity.

Mutation or deletion of chTOG appeared to stabilize the erroneous attachments generated by STLC treatment, so we next directly tested whether chTOG is required to *correct* errors. We washed out the STLC and assayed if cells recovered from the monopolar state and aligned chromosomes. Immediately after STLC removal, few cells had aligned chromosomes in any experimental condition (Figure 4 – Figure Supplement 1b). 60 minutes after STLC washout, a majority of control cells (not treated with siRNA) had formed a bipolar spindle and aligned chromosomes at the spindle equator (Fig. 4d). As expected, chTOG depleted cells failed to form bipolar spindles and therefore did not align chromosomes at a metaphase plate (Fig. 4d) (Cassimeris & Morabito, 2004; Gergely et al., 2003). In contrast, expression of either chTOG^WT^ or chTOG^KK/AA^ after siRNA treatment rescued bipolar spindle formation after 60 minutes; however, only chTOG^WT^ expressing cells properly aligned their chromosomes (Fig. 4d). Together, these data reveal that chTOG functions as a mitotic error correction factor, and this activity appears independent of its well-characterized role as a regulator of the microtubule cytoskeleton.

**Figure 4 – Figure Supplement 1.**
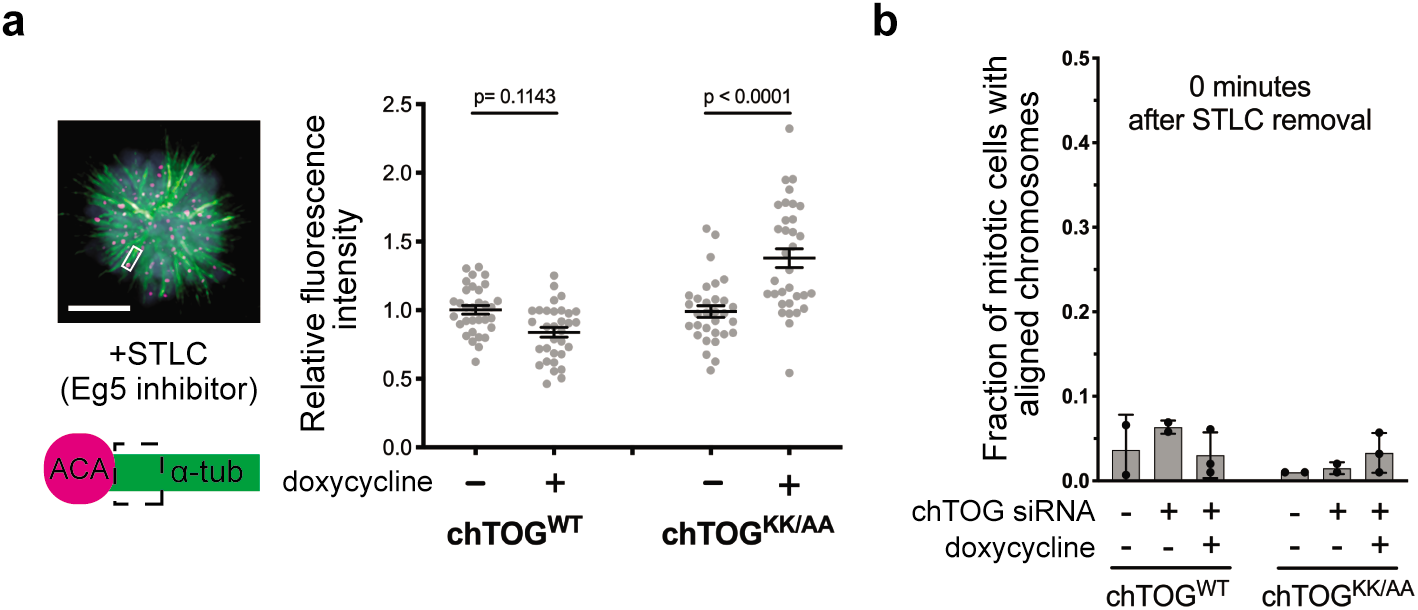
The basic pair mutant prematurely stabilizes erroneous kinetochore microtubule attachments even in the presence of endogenous chTOG. (a) HeLa cells expressing chTOG™ and chTOG^KK/AA^ in the presence of endogenous chTOG were arrested with STLC to form monopolar spindles and erroneous low-tension kinetochore-microtubule attachments. Immunofluorescence was performed with anti-tubulin and anti-ACA antibodies and the intensity of tubulin staining was quantified near kinetochores as an indirect measure of the number of microtubules within an attached bundle. The basic pair mutant prematurely stabilizes erroneous attachments even in the presence of endogenous chTOG. (b) Cells under all experimental conditions in Figure 4 were quantified for the fraction of mitotic cells with aligned chromosome immediately after STLC removal. In every condition, cells responded equivalently to STLC treatment and washout.

### chTOG-based error correction is independent of Hec1 phospho-regulation

Our work suggested that chTOG functions similarly to the budding yeast ortholog (Stu2) that confers an intrinsic tension-dependent microtubule binding behavior to kinetochores that is independent of the extrinsic signaling through the Aurora B pathway (Miller et al., 2016, 2019). To test this in human cells, we analyzed recovery from STLC when both pathways were inhibited. Because the complete inhibition of Aurora B with ZM447439 (ZM) prevents cells from forming bipolar spindles after STLC washout (K. F. DeLuca, Lens, & DeLuca, 2011) (data not shown), we partially inhibited Aurora B with a lower dose. Cells expressing WT chTOG while Aurora B was inhibited exhibited the same phenotype as ZM treatment alone. However, when the basic pair mutant was expressed in the presence of ZM, we observed an additive phenotype where essentially no cells formed an organized metaphase plate after 60 minutes (Fig. 5a). These data suggested that chTOG does not regulate the Aurora B pathway. To further examine this, we measured the phosphorylation status of the Aurora B substrate Hec1 upon chTOG depletion. We assayed cells containing unaligned chromosomes and found that chTOG depletion does not affect the phosphorylation status of Ser55 on Hec1, while partial inhibition of Aurora reduced the prevalence of this mark by ~50% (Fig. 5b). These findings strongly suggest that the chTOG error correction pathway is independent of Aurora B mediated error correction.

**Figure 5.**
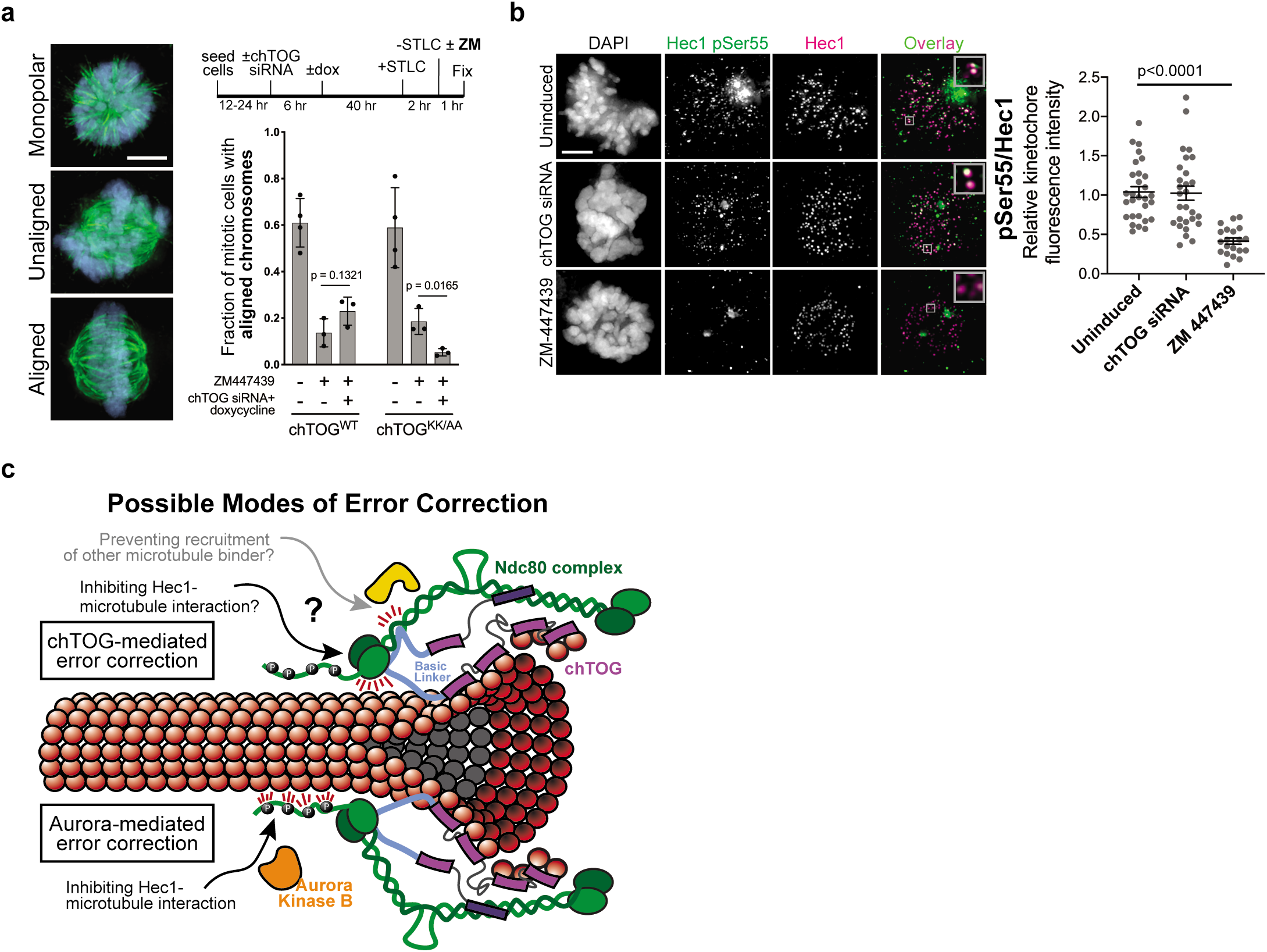
chTOG- and Aurora B-dependent error pathways likely function independently. (a) The same mitotic error correction assay from Fig. 4d was performed in the presence of Aurora B inhibitor ZM447439. The mean and standard deviation of three experimental replicates are displayed, (b) Representative immunofluorescence images (left) and quantifications (right) of the relative fluorescence intensity of phosphorylated Hec1 analyzed with a phospho-specific antibody to Ser55. Each data point represents the average kinetochore intensity within a single cell normalized to the mean value of mock transfected cells. The relative intensity was determined for both Hec1 pSer55 and Hec1 antibodies, then ratioed. Mean and standard error of the mean for three experimental replicates displayed. All p values were determined by Tukey’s multiple comparisons test, and all error bars are 5 μm. (c) Possible models of chTOG-mediated error correction (top) where chTOG kinetochore localization depends on the C-terminus while the basic linker inhibits Ndc80 complex activity. We favor the basic linker directly modulating Hec1 microtubule binding behavior, but chTOG could also prevent recruitment of other microtubule binding proteins. This activity is independent of Aurora-mediated error correction (bottom) phosphorylating the Ndc80 tail domain to destabilize attachments.

## Discussion

Here we have identified a previously unknown mitotic error correction pathway in human cells, where chTOG localizes to kinetochores and destabilizes erroneous, low-tension attachments. While numerous proteins have been implicated in selectively stabilizing bioriented attachments (Gaitanos et al., 2009; Girão et al., 2020; Janke, Ortíz, Tanaka, Lechner, & Schiebel, 2002; Maiato et al., 2003; Ortiz, Funk, Schäfer, & Lechner, 2009; Raaijmakers, Tanenbaum, Maia, & Medema, 2009; Schmidt et al., 2010; Welburn et al., 2009), less is known about the mechanisms that recognize and destabilize errors. This activity of chTOG was likely not previously elucidated due to the pleiotropic effects of chTOG depletion on spindle structure and dynamics. By arresting cells with monopolar spindles and identifying a targeted mutation of the basic linker, we were able to specifically assay chTOG error correction functions independent of its regulation of the cytoskeleton. Although the basic linker regulates polymerase activity *in vitro* via non-specific electrostatic interactions with the microtubule lattice (Geyer et al., 2018; Wang & Huffaker, 1997; Widlund et al., 2011), yeast viability was normal when the entire basic linker (pI = 10.4; 18 positive residues) was replaced with just 15 conserved residues (pI = 8.7; 3 positive residues). Moreover, mutation of a single positive residue rendered cells completely inviable. Therefore, while the basic linker net charge is a determinate for *in vitro* polymerase activity (Geyer et al., 2018; Widlund et al., 2011), it does not correlate with cell viability. We therefore propose that the essential function of the basic linker is to mediate error correction rather than regulate microtubule dynamics.

We also found that the conserved process of chTOG-based error correction is most likely independent of Aurora B phosphorylation of Hec1, similar to budding yeast (Miller et al., 2019). In fact, our data suggest that Aurora B activity at the kinetochore does not respond to or compensate for defects in chTOG-mediated error correction. Thus, we envision two error correction pathways that respond to separate input signals and leverage unique molecular strategies to destabilize erroneous kinetochore-microtubule attachments. Why do cells utilize multiple mechanisms of error correction? One possibility is the intrinsic tension-sensing pathway uses chTOG to ‘read’ the structure and position of the microtubule tip and respond to tension across the kinetochore-microtubule bond. Meanwhile, the Aurora B-mediated extrinsic pathway is instead regulated by tension transmitted through the inner-kinetochore and centromere to regulate the balance of kinase and phosphatase activity on microtubule binding proteins. Thus, each pathway would respond to unique but overlapping tension-dependent changes that occur as kinetochore-microtubule attachments mature. Aurora B does not appear to compensate for the loss of chTOG-based error correction, further suggesting these pathways respond to unique stimuli and cells utilize each mechanism to fine tune their response to forces experienced during attachment.

One similarity between the two pathways is they converge at the Ndc80 complex. While the exact mechanism remains unclear, chTOG likely inhibits Hec1 to destabilize erroneous attachments rather than by directly modulating microtubules. We suggest this because the basic pair mutant retains the ability to bind to kinetochores and microtubule tips, yet it fails to destabilize erroneous kinetochore-microtubule attachments similar to depletion of chTOG (Fig. 5c). Thus, we favor a model where the C-terminus of Stu2/chTOG is required for stable association with the kinetochore (Miller et al., 2019) while the basic linker modulates Hec1’s microtubule binding behavior through competitive or allosteric means (Fig. 5c). While it is also possible the basic linker inhibits the recruitment of other microtubule binding factors to the Nc80 complex, this is less likely because most of these factors are unique to budding yeast (Dam1/DASH complex) or metazoans (SKA and Astrin-SKAP complexes) (Hooff, Kops, Tromer, & Wijk, 2017; Van Hooff, Snel, & Kops, 2017), yet the basic linker functions similarly in both species. Understanding the relationship between the Ndc80 complex and the basic linker in the context of microtubule attachment will be a key area of future research.

chTOG error correction activity also provides new perspectives for cancer biology. RNAi screens for cancer vulnerabilities suggested that some tumors grow dependent or ‘addicted’ to elevated levels of chTOG, making it a therapeutic target. However, chTOG depletion is reminiscent of microtubule poisons that are universally toxic, but particularly potent in specific cancers (Martens-de Kemp et al., 2013; Tiedemann et al., 2012). Dose limiting toxicities are a common challenge for developing anti-mitotic therapies, but preclinical studies indicate that they can be overcome through precision inhibition of single functions within multifunctional proteins (Ding et al., 2013). Future analyses of the basic pair mutant will reveal if specifically inhibiting chTOG-mediated error correction, rather than degrading the entire protein, is therapeutically advantageous.

## Methods

### Mammalian cell culture

HCT116 (Cherry et al., 2019), 293T (Ding et al., 2013), and HeLa FlpIn Cells (Etemad, Kuijt, & Kops, 2015) cells were grown in a high glucose DMEM (Thermo Fisher Scientific 11-965-118/Gibco 11965118) supplemented with anti/anti (Thermo Fisher Scientific 15240062) and 10% Foetal Bovine Serum (Thermo Fisher Scientific 26140095) at 37 °C supplemented with 5% CO2. For microscopy experiments, cell suspensions were added to 35mm wells containing acid washed 1.5 x 22mm square coverslips (Fisher Scientific 152222) and grown for 12-24 hours prior to transfections or immunostaining. For live cell microscopy experiments, cell suspensions were added to 35 mm glass bottom dishes (Mattek Corp. P35G-1.5-20-C) and grown in standard growth media because they were performed in an environmental chamber using TIRF microscopy.

To entirely depolymerize the microtubule cytoskeleton, growth media were supplemented with 10 μM nocodazole (Sigma-Aldrich, M1404) for one hour. To synchronize cells, they were treated with 2.5 mM thymidine (Sigma-Aldrich, T9250) for 16 hours, cells were then placed in drug-free media for 8 hours. A second synchronization was achieved by supplementing media with 2.5 mM thymidine again for 16 hours. Finally, thymidine was removed and 4-5 hours later (depending on cell line) 10 μM nocodazole was added. Eg5 inhibition was achieved by supplementing growth media with 5 μM S-trityl-L-cysteine (STLC, Sigma-Aldrich, 164739) for 2 hours. Partial inhibition of Aurora B kinase was performed with a one-hour long treatment of 500 nM ZM447439 (Selleckchem, S1103).

### Immunoprecipitations

15 cm dishes of 293T cells were transfected with 40 μg of empty vector (pSB2353), chTOG^WT^-3Flag (pSB2976), or chTOG^KK/AA^-3Flag (pSB2977) using 85 μg of polyethyleneimine (PEI, Polysciences 23966-1) as previously reported (Longo, Kavran, Kim, & Leahy, 2013). Cells were harvested by mechanical dissociation with PBS and then centrifuged. The cell pellet was resuspended in 250 μL of complete lysis buffer [25mM HEPES, 2 mM MgCl2, 0.1 mM EDTA, 0.5 mM EGTA, 15% Glycerol, 0.1% NP-40, 150 mM KCl, 1 mM PMSF, 1 mM sodium pyrophosphate, 1x Pierce Protease Inhibitor Cocktail (Thermo Scientific 88666)] and snap frozen in liquid nitrogen. Samples were thawed and sonicated for 20 seconds at 50% intensity three times. Approximately 150 U of Benzonase nuclease (Millipore E1014) was added to samples and incubated at room temperature for five minutes. The samples were centrifuged at maximum speed at 4 °C in a tabletop centrifuge. Clarified lysates were moved to fresh microfuge tubes and 60 μL of protein A dynabeads (Thermo Fisher 10001D) conjugated with anti-FLAG(M2) monoclonal antibody (Sigma Aldrich F3165) as previously described (Akiyoshi, Nelson, Ranish, & Biggins, 2009) were added and incubated at 4 °C with rotation for 90 minutes. Beads were washed four times with lysis buffer lacking PMSF, sodium pyrophosphate, and protease inhibitor cocktail. Proteins were eluted from beads in 40 μL of SDS sample buffer incubated at 95 °C for 5 minutes.

### Immunoblotting

Clarified lysates prepared as described above were diluted in 2x SDS sample buffer and incubated at 95 °C for 5 minutes. Immunoblots for chTOG were prepared by resolving lysates on NuPAGE 4-12% Bis-Tris Gels (Life Technologies, NP0329BOX) in 1x MOPS-SDS buffer and then transferring the proteins to 0.45 μm nitrocellulose membrane (BioRad, 1620115) for two hours at 4 °C in transfer buffer containing 20% methanol. Membranes were washed in PBS+0.05% Tween-20 (PBS-T) and blocked with PBS-T+5% non-fat milk overnight at 4 °C. Primary antibodies were diluted in PBS-T by the following factors or to the following concentrations: anti-GAPDH clone 6C5 (Millipore Sigma, MAB374) 1 μg/mL; anti-CKAP5(chTOG) (GeneTex, GTX30693) 1:1000; anti-HEC1 clone 9G3 (ThermoFisher Scientific MA1-23308) 2 μg/mL. HRP conjugated antimouse (GE Lifesciences, NA931) and anti-rabbit (GE Lifesciences, NA934) secondary antibodies were diluted 1:10,000 in PBS-T and incubated on membranes for 45 minutes at room temperature. Immunoblots were developed with enhanced chemiluminescence HRP substrates SuperSignal West Dura (Thermo Scientific, 34076) or SuperSignal West Femto (Thermo Scientific, 34094). All chemiluminescence was detected using a ChemiDoc MP system (BioRad).

### Immunofluorescent Staining

Upon completion of experimental manipulations, cells grown on coverslips were immediately chemically crosslinked for 15 minutes with 4% PFA diluted from a 16% stock solution (Electron Microscopy Sciences, 15710) with 1x PHEM (60 mM PIPES, 25 mM HEPES, 5 mM EGTA, 8 mM MgSO4). The exception was experiments where HEC1 levels were quantified, in which cells were treated with 1x PHEM+0.5% TritonX100 for 5 minutes prior to PFA. Coverslips were washed with 1x PHEM+0.5% TritonX100 for 5 minutes, then washed 3 more times with 1x PHEM + 0.1% TritonX100 over 10 minutes. Cells were blocked for 1-2 hours at room temperature in 20% goat serum in 1x PHEM. Primary antibodies were diluted in 20% goat serum to the following final concentrations / dilution factors: anti-centromere protein antibody or ACA (Antibodies Inc. 15-235) 1:600; anti-HEC1 clone 9G3 (ThermoFisher Scientific, MA1-23308) 2 μg/mL; anti-alpha tubulin clone DM1A (Sigma Millipore, T6199) 2 μg/mL; anti-Mad1 (GeneTex, GTX109519) 1:1000, anti-Hec1pSer55 (Jennifer DeLuca, Colorado State University) 1:1000). Coverslips were incubated overnight at 4°C in primary antibody, then washed four times with 1x PHEM + 0.1% TritonX100 over 10 minutes. Goat anti-mouse, rabbit, and human secondary antibodies conjugated to AlexaFluor 488, 568, 647 (Invitrogen) were all diluted at 1:300 in 20% boiled goat serum with the exception of goat anti-mouse AlexaFluor647 used to target mouse anti-alpha tubulin where 1:600 dilution was used. Coverslips were washed four times with 1x PHEM + 0.1% TritonX100 over 10 minutes, then stained for one minute with 30 ng/mL 4’,6-diamidino-2-phenylindole (DAPI, Invitrogen, D1306) in 1x PHEM. Coverslips were washed two times with 1x PHEM, then immersed in mounting media (90% glycerol, 20 mM Tris [pH= 8.0], 0.5% w/v N-propyl gallate) on microscope slides and sealed with nail polish.

### Microscopy and Image Analysis

Fixed cell images were acquired on either a Deltavision Elite or Deltavision Ultra deconvolution high-resolution microscope, both equipped with a 60x/1.42 PlanApo N oil-immersion objective (Olympus). Slides imaged on the Elite were collected with a Photometrics HQ2 CCD 12-bit camera, while those imaged on the Ultra were equipped with a 16-bit sCMOS detector. On both microscopes, cells were imaged in Z-stacks through the entire cell using 0.2 μm steps. All images were deconvolved using standard settings. Softworx Explorer 2.0 was used to quantify kinetochore intensities by identifying the maximal ACA intensity within a Z-stack and collecting pixel intensity with a 16-pixel region of interest for the appropriate wavelength, as well as a 36-pixel region encompassing the first region for background subtraction. Background intensity was found by subtracting the intensity of the 16-pixel region from the 36-pixel region. This background intensity was then divided by the difference in area (20 pixels) to give background intensity per pixel. This was then multiplied by 16 and subtracted from the initial intensity of the 16-pixel region. Representative images displayed from these experiments are projections of the maximum pixel intensity across all Z images. Photoshop was used to crop, make equivalent, linear adjustments to brightness and contrast, and overlap images.

Live cell TIRF microscopy was performed on a Nikon widefield fluorescence and TIRF microscope equipped with an 100x/1.49 CFI Apo TIRF oil immersion objective (Nikon) and an Andor iXon X3 EMCCD camera. Cells were imaged briefly with dual lasers to visualize expression of EB1-mCherry and when appropriate, chTOG-EGFP. TIRF images for quantification were collected every 300 ms over a 90 second period using only the 561 nm laser to monitor EB1-mCherry. Tiff files for each timepoint were imported into FIJI (Schindelin et al., 2012), background subtracted, and ROF denoised prior to semi-automated track analysis with the TrackMate plugin (Tinevez et al., 2017) using the DoG Detector and Simple LAP Tracker. Kymographs generated from background subtracted movies with Fiji (Schindelin et al., 2012) using a 7 μm line profile over the entire duration of the experiment.

### Statistics

GraphPad Prism version 8.4 was used for statistical analysis. Data normality was assessed for all experiments using Shapiro-Wilk, Anderson-Darling, D’Agostino & Pearson, and Kolmogorov-Smirnov tests. For those with normal distributions, mean values were reported and appropriate statistical comparison tests were used. For data sets failing normality tests, median values were reported, and non-parametric comparisons were performed. Each test specifically identified in figure legends.

### Multiple Sequence Alignments

Fungal proteins related to *S. cerevisiae* Stu2 were identified using a PSI-BLAST (Altschul et al., 1997) search on NCBI. Multiple sequence alignments of the entire proteins were generated with ClustalOmega (Sievers et al., 2011) default parameters and displayed in JalView 1.8(Waterhouse, Procter, Martin, Clamp, & Barton, 2009). Eukaryotic basic linker sequences were manually identified and aligned with ClustalOmega for display in JalView.

### Nucleic Acid Reagents

All plasmids used in this study are described in Supplementary Table 1. Construction of a *LEU2* integrating plasmid containing wild-type *pSTU2-STU2-3V5* (pSB2232) is described before (Miller et al., 2016). *STU2* variants were constructed by mutagenizing pSB2232 as described previously (H. Liu & Naismith, 2008; Tseng, Lin, Wei, & Fang, 2008), resulting in pSB2634 (*pSTU2-stu2(Δ592-607::GDGAGL^linker^)-3V5*, i.e. *stu2^ΔPatch^)*, pSB2781 (*pSTU2-stu2(K598A R599A)-3V5*, i.e. *stu2^KR/AA^)*, pSB2818 (*pSTU2-stu2(K598A)-3V5*, i.e. *stu2^K598A^)*, pSB2819 (*pSTU2-stu2(R599A)-3V5*, i.e. *stu2^R599A^)*, pSB2260 (Miller et al., 2019) (*pSTU2-stu2(Δ560-657::GDGAGL^linker^)-3V5*, i.e. *stu2^ΔBL^)*, pSB2820 (*pSTU2-stu2(Δ551-657::GDGAGL^linker^:592-607:GDGAGL^linker^)-3V5)*. pSB2820 was further mutagenized following the above protocols resulting in pSB2852 (*pSTU2-stu2(Δ560-657::GDGAGL^linker^:592-607:GDGAGL^linker^)-3V5*, i.e. *stu2^ΔBL+Patch^*).

Codon-optimized chTOG^WT^ and chTOG^KK/AA^ were synthesized and sub-cloned into pCDNA3.1-C-EGFP by Genscript (pSB2822 and pSB2823, respectively). Both chTOG^WT^ and chTOG^KK/AA^-EGFP fusions were cloned into pCDNA5 FRT/TO/puro (pSB2353) (Etemad et al., 2015) through PCR amplification and isothermal assembly to generate pSB2860 and pSB2863, respectively. EGFP was excised from pSB2860 and pSB2863 through restriction digestion and replaced with 6-His,3-FLAG via isothermal assembly to generate pSB2976 and pSB2977. mCherry-EB1 fusion was PCR amplified from mCherry-EB1-8, a gift from Michael Davidson (Addgene plasmid # 55035), and inserted in third generation lentiviral vector, pLPH2 (pSB2998), via isothermal assembly to generate pSB3217. Primers used in the construction of the above plasmids are listed in Supplementary Table 1, further details of plasmid construction including plasmid maps available upon request.

The siRNA targeting chTOG and Hec1 were ordered from Qiagen. Hec1 was depleted with a custom synthesized siRNA sequence (5’-CCCUGGGUCGUGUCAGGAA-3’) that targets the 5’ UTR and was previously validated (K. F. DeLuca et al., 2011). Hs_ch-TOG_6 FlexiTube siRNA (SI02653588) targets the coding DNA sequence of endogenous chTOG but not the codon optimized EGFP fusion.

### Generation of Yeast Strains

*Saccharomyces cerevisiae* strains used in this study are described in Supplementary Table 2 and are derivatives of SBY3 (W303). Standard media, microbial, and genetic techniques were used(Sherman, Fink, & Lawrence, 1974). Stu2-3HA-IAA7 was constructed by PCR-based methods (Longtine et al., 1998) and is described previously (Miller et al., 2016).

### Yeast Growth Assay

The desired strains were grown overnight in yeast extract peptone plus 2% glucose (YPD) medium. The following day, cells were diluted to OD600 ~1.0 from which a serial 1:5 dilution series was made and spotted on YPD+DMSO or YPD+100 μM IAA (indole-3-acetic acid dissolved in DMSO) plates. Plates were incubated at 23°C for 3 days.

### Generation of Modified Human Cell Lines

All human cell lines used in this study are described in Supplementary Table 3. 400,000 HeLa FlpIn TREX cells (SBM001) were grown in 60mm dishes for 16 hours. Media was aspirated and replaced with 2.5 mL of DMEM containing no supplements. 3.2 μg of p0G44 (pSB2380) and 1 μg of pSB2860/pSB2863 were mixed with 8 μL of Lipofectamine2000 (Invitrogen 11668027) in 400 μL of DMEM (no supplements) for 20 minutes and then added to cells dropwise. Six hours after transfection, media was aspirated and replaced with DMEM containing 10% FBS and antibiotics. 48 hours post transfection, media was supplemented with 2.5 μg/mL puromycin and cells were negatively selected for 3 days. Upon reaching confluence, expression of EGFP fusion proteins was induced by addition of 1 μg/mL doxycycline (Sigma-Aldrich, D9891) and EGFP expressing cells were positively selected by FACS using a SONY MA900 to sort into media lacking doxycycline. Doubly selected polyclonal populations of chTOG^WT^-EGFP (SBM044) and chTOG^KK/AA^-EGFP (SBM046) were frozen and stored for future experiments.

EB1-mCherry was stably transduced into SBM044 and SBM046 via lentivirus. Assembly of replication deficient viral particles was performed as previously described (Toledo et al., 2014). Briefly, pLCH2-EB1-mCherry (pSB3217), pPAX2 (pSB2636), and pMD2.G (pSB2637) were cotransfected into 293T cells using PEI. Virus containing supernatant media were harvested 48 hours post transfection and passed through 0.45 μm filters and frozen at −80°C. Filtered viral supernatant media were added to dishes containing SBM044 and SBM046 and 48 hours later hygromycin B (Invitrogen 10687010) was added at 400 μg/mL. Cells were selected for 5 days to generate SBM045 and SBM047.

## Acknowledgments

We thank Linda Wordeman, Mike Wagenbach, and Juan-Jesus Vicente for HCT116 cells with chTOG endogenously EGFP tagged and help with EB1-tracking; Geert Kops for HeLa FlpIn TREX cells and pCDNA5 FRT/TO vectors; Jennifer DeLuca for Hec1 antibodies; Michael Davidson for EB1-mCherry DNA; the entire Biggins Lab and Chip Asbury for feedback on this manuscript.

## Supplementary Note 1

MT assembly rates have been reported to decrease after chTOG depletion by RNAi in some studies (Ertych et al., 2014; van der Vaart et al., 2011), while no changes were observed in a different study (Cassimeris et al., 2009). In our work, we observed increased MT assembly rates. We believe these differences arise from our use of lentivirus to stably express EB1-mCherry from the human PGK1 promoter, while all three other studies transiently over-expressed the EB protein from a highly active CMV promoter. While cells tolerate EB1 over-expression relatively well, high levels of expression and the length of the linkage between EB1 and EGFP have both been shown to alter MT polymerization rates (Geisterfer, Zhu, Mitchison, Oakey, & Gatlin, 2020; Skube, Chaverri, & Goodson, 2010). Because of these differences we believe the increase in polymerization rates we observe is a result of changes in the cellular pool of free tubulin that increases because chTOG no longer contributes to MT nucleation or the sequestration of free dimers. In addition, altered tubulin homeostasis has been observed with siRNA depletion of numerous kinesins (Wordeman, Decarreau, Vicente, & Wagenbach, 2016) and could contribute to our results.

## Supplementary Tables

**Supplementary Table 1.** Plasmids used in this study.

| Plasmid | Vector Backbone | Gene of Interest | Mutation Description | Selection Marker | Primers | Source |
| --- | --- | --- | --- | --- | --- | --- |
| pSB2232 | pSB2223 / pL300 | Stu2<sup>WT</sup>-3V5 |  | LEU2 | SB4372, SB4374 | Miller et al., 2016 |
| pSB2260 | pSB2223 / pL300 | Stu2<sup>ΔBL</sup>-3V5 | Δ560-657 :: GDGAG | LEU2 | SB4411, SB4413 | Miller et al., 2019 |
| pSB2634 | pSB2223 / pL300 | Stu2<sup>ΔPatch</sup>-3V5 | Δ592-607 :: GDGAG | LEU2 | SB5248, SB4413 | This study |
| pSB2820 | pSB2223 / pL300 | Cloning intermediate | Δ569-657:: GDGAG+592-607+GDGAG | LEU2 | SB5447 | This study |
| pSB2852 | pSB2223 / pL300 | Stu2<sup>ΔBL+Patch</sup>-3V5 | Δ560-657:: GDGAG+592-607+GDGAG | LEU2 | SB5519, SB5520 | This study |
| pSB2818 | pSB2223 / pL300 | Stu2<sup>ΔK598A</sup>-3V5 | K598A | LEU2 | SB5458, SB4413 | This study |
| pSB2819 | pSB2223 / pL300 | Stu2<sup>ΔR599A</sup>-3V5 | R599A | LEU2 | SB5459, SB4413 | This study |
| pSB2781 | pSB2223 / pL300 | Stu2<sup>ΔKR598AA</sup>-3V5 | K598A, R599A | LEU2 | SB5349, SB4413 | This study |
| pSB2822 | pCDNA3.1 | chTOG<sup>WT</sup>-EGFP |  | Neomycin | N/A | This study |
| pSB2823 | pCDNA3.1 | chTOG<sup>KK1142AA</sup>-EGFP | K1142A, K1143A | Neomycin | N/A | This study |
| pSB2353 | pCDNA5 FRT/TO | N/A |  | Puromycin | N/A | Etemad et al., 2015 |
| pSB2860 | pCDNA5 FRT/TO | chTOG<sup>WT</sup>-EGFP |  | Puromycin | SB5536, SB5537 | This Study |
| pSB2863 | pCDNA5 FRT/TO | chTOG<sup>KK1142AA</sup>-EGFP | K1142A, K1143A | Puromycin | SB5536, SB5537 | This Study |
| pSB2976 | pCDNA5 FRT/TO | chTOG<sup>WT</sup>-6His-3Flag |  | Puromycin | SB5774, SB5775 | This Study |
| pSB2977 | pCDNA5 FRT/TO | chTOG<sup>KK1142AA</sup>-6His-3Flag | K1142A, K1143A | Puromycin | SB5774, SB5775 | This Study |
| pSB3062 |  | EB1-mCherry |  | Neomycin | N/A | Davidson Lab (Addgene: 55035) |
| pSB2998 | pLPH2 | N/A |  | Hygromycin | N/A | This Study |
| pSB3217 | pLPH2 | EB1-mCherry |  | Hygromycin | SB5938, SB5939, SB5940, SB5941 | This Study |

**Supplementary Table 2.** Yeast strains used in this study.

All strains are derivatives of SBY3 (W303)
| Strain | Relevant Genotype |
| --- | --- |
| SBY3 (W303) | *MATa ura3-1 leu2-3,112 his3-11 trp1-1 can1-100 ade2-1 bar1-1* |
| SBY13772 | *MATa STU2-3HA-IAA7:KanMX DSN1-6His-3Flag:URA3 his3::pGPD1-TIR1:HIS3* |
| SBY13901 | *MATa STU2-3HA-IAA7:KanMX DSN1-6His-3Flag:URA3 his3::pGPD1-TIR1:HIS3 leu2::pSTU2-STU2-3V5:LEU2* |
| SBY17069 | *MATa STU2-3HA-IAA7:KanMX DSN1-6His-3Flag:URA3 his3::pGPD1-TIR1:HIS3 leu2::pSTU2-stu2(Δ592-607::GDGAGLinker)-3V5:LEU2* |
| SBY17206 | *MATa STU2-3HA-IAA7:KanMX DSN1-6His-3Flag:URA3 his3::pGPD1-TIR1:HIS3 leu2::pSTU2-stu2(K598A R599A)-3V5:LEU2* |
| SBY17477 | *MATa STU2-3HA-IAA7:KanMX DSN1-6His-3Flag:URA3 his3::pGPD1-TIR1:HIS3 leu2::pSTU2-stu2(K598A)-3V5:LEU2* |
| SBY17479 | *MATa STU2-3HA-IAA7:KanMX DSN1-6His-3Flag:URA3 his3::pGPD1-TIR1:HIS3 leu2::pSTU2-stu2(R599A)-3V5:LEU2* |
| SBY17519 | *MATa STU2-3HA-IAA7:KanMX DSN1-6His-3Flag:URA3 his3::pGPD1-TIR1:HIS3 leu2::pSTU2-stu2(Δ560-657::GDGAGLinker)-3V5:LEU2* |
| SBY17593 | *MATa STU2-3HA-IAA7:KanMX DSN1-6His-3Flag:URA3 his3::pGPD1-TIR1:HIS3 leu2::pSTU2-stu2(Δ560-657::GDGAGLinker:592-607:GDGAGLinker)-3V5:LEU2* |

**Supplementary Table 3.**
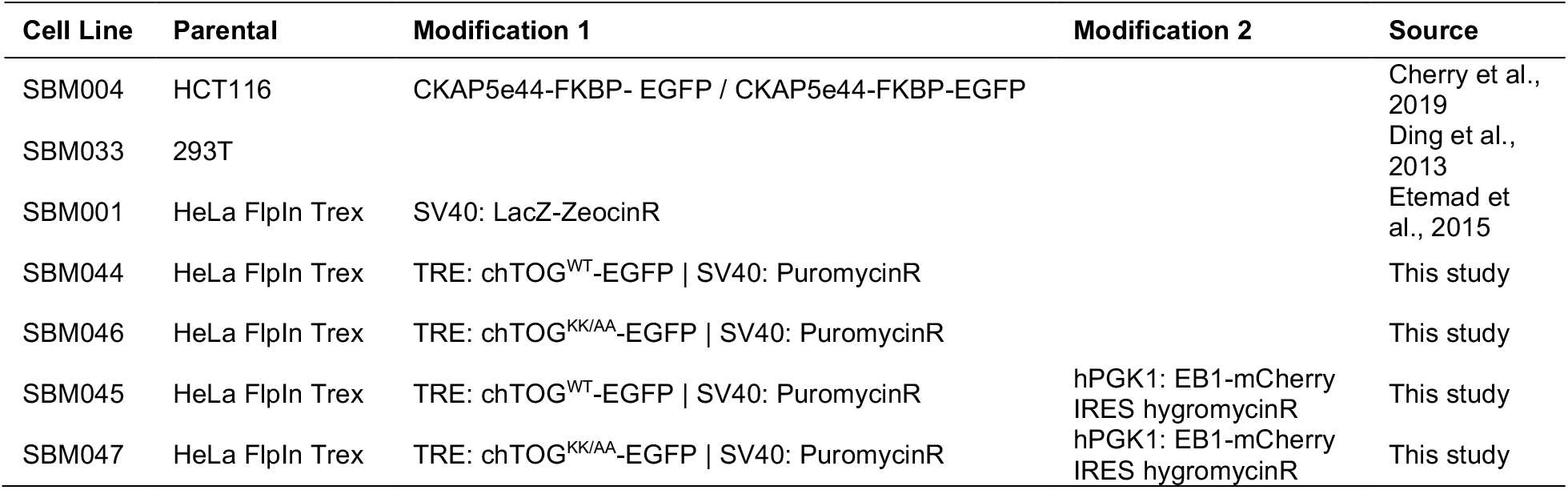
Human cell lines used in this study.

| Cell Line | Parental | Modification 1 | Modification 2 | Source |
| --- | --- | --- | --- | --- |
| SBM004 | HCT116 | CKAP5e44-FKBP- EGFP / CKAP5e44-FKBP-EGFP |  | Cherry et al., 2019 |
| SBM033 | 293T |  |  | Ding et al., 2013 |
| SBM001 | HeLa FlpIn Trex | SV40: LacZ-ZeocinR |  | Etemad et al., 2015 |
| SBM044 | HeLa FlpIn Trex | TRE: chTOG<sup>WT</sup>-EGFP \| SV40: PuromycinR |  | This study |
| SBM046 | HeLa FlpIn Trex | TRE: chTOG<sup>KK/AA</sup>-EGFP \| SV40: PuromycinR |  | This study |
| SBM045 | HeLa FlpIn Trex | TRE: chTOG<sup>WT</sup>-EGFP \| SV40: PuromycinR | hPGK1: EB1-mCherry<br>IRES hygromycinR | This study |
| SBM047 | HeLa FlpIn Trex | TRE: chTOG<sup>KK/AA</sup>-EGFP \| SV40: PuromycinR | hPGK1: EB1-mCherry<br>IRES hygromycinR | This study |

